# Chemo-sEVs release in cisplatin-resistance ovarian cancer cells are regulated by the lysosomal function

**DOI:** 10.1101/2023.02.03.526974

**Authors:** Cristóbal Cerda-Troncoso, Felipe Grünenwald, Eloísa Arias-Muñoz, Viviana A. Cavieres, Albano Caceres-Verschae, Sergio Hernández, Belén Gaete-Ramírez, Francisca Álvarez-Astudillo, Rodrigo A Acuña, Matias Ostrowski, Patricia V. Burgos, Manuel Varas-Godoy

## Abstract

Ovarian cancer (OvCa) is an aggressive disease usually treated with cisplatin (CDDP)-based therapy. However, among the different types of cancers treated with CDDP, OvCa commonly develops chemoresistance to this treatment. The small extracellular vesicles (sEVs) play a central role in chemoresistance. In response to chemotherapy, resistant cells secrete sEVs named chemo-sEVs characterized by specific cargo landscape content involved in the transfer of chemoresistance to recipient cells. sEVs encompass a variety of vesicle types, including exosomes, and are formed as intraluminal vesicles (ILVs) within multivesicular endosomes (MVEs). MVEs follow at least two trafficking pathways regulated by RAB GTPase family members; 1) a secretory pathway where MVEs fuse with the plasma membrane (PM) for sEVs secretion, where RAB27A is the most studied; 2) a degradative pathway where MVEs fuse with lysosomes, an event controlled by RAB7. There is growing evidence suggesting that a loss of lysosomal function can increase sEVs secretion; however, whether sEVs secretion and the transfer of CDDP chemoresistance in OvCa is the result of a fine regulation between these two MVEs trafficking pathways is unknown. In this work, we study the status of these two pathways, between CDDP-sensitive (A2780) and CDDP-resistant (A2780cis) OvCa cells. We found A2780cis cells have an increased number of MVEs and ILVs structures, together with higher levels of ESCRTs machinery components and RAB27A, compared to A2780 cells. Moreover, CDDP promotes the secretion of chemo-sEVs in A2780cis cells. Interestingly, chemo-sEVs contain a high number of proteins related to DNA damage response. In addition, we determine A2780cis cells have a poor lysosomal function with reduced levels of RAB7. Surprisingly, silencing of RAB27A in A2780cis cells was found to be sufficient to restore lysosomal function and levels of RAB7 in A2780cis cells, switching into an A2780-like cellular phenotype. Next, we found rapamycin, a potent enhancer of lysosomal function, reduced the secretion of chemo-sEVs. Taken together, these results indicate that the secretion of chemo-sEVs in OvCa cells is determined by the balance between secretory MVEs and MVEs that are destined for lysosomal degradation. Thus, our results suggest that adjusting this balance between these two MVEs trafficking pathways could be a promising strategy for overcoming CDDP chemoresistance in OvCa.

**Graphical Abstract:** 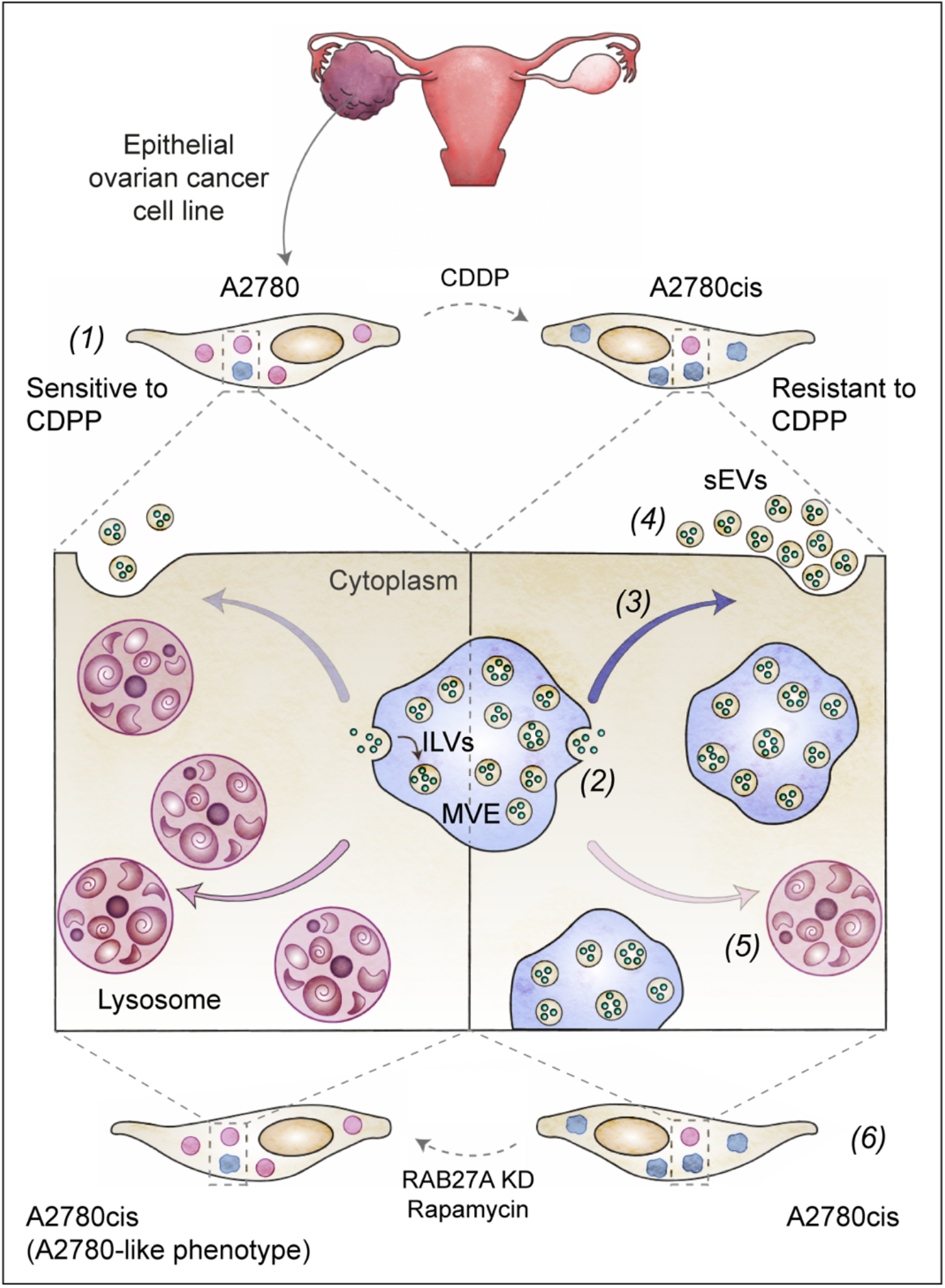

## Introduction

The World Health Organization reported that around 18.1 million women worldwide had cancer, and 9.6 million died due to this disease (Bray et al., 2018). Ovarian cancer (OvCa) is one of the most common gynecologic cancers (Sung et al., 2021) The current treatment of OvCa involves primary debulking surgery followed by platinum-based and/or taxane-based combination chemotherapy (Kim et al., 2018; Reid et al., 2017). For platinum-based, cisplatin (CDDP) is the most used chemotherapy (Dasari & Bernard Tchounwou, 2014; Helm & States, 2009), however a significant percentage of sensitive cells develop chemoresistance (Galluzzi et al., 2012; Shen et al., 2012). Moreover, the cellular processes responsible for CDDP chemoresistance in OvCa are poorly understood.

The acquisition of chemoresistance depends on adaptive responses of the tumor cells to evade the effects of chemotherapy (Qin et al., 2020; Tchounwou et al., 2021). In OvCa, CDDP-chemoresistance includes a reduction in the entry of CDDP inside of cells, DNA repair mechanisms, metabolic reprogramming, cell death inhibition; and epigenetic changes (Ai et al., 2016; Chan et al., 2021; Norouzi-Barough et al., 2018; Patch et al., 2015; Tchounwou et al., 2021; Tong et al., 2019; Xie et al., 2021). In addition, there is an important contribution of the small extracellular vesicles (sEVs) that participate in intercellular communication through the secretion of proteins and genetic material (Mathieu et al., 2021). Importantly, different models of chemoresistant cells secrete chemo-sEVs in response to chemotherapy including OvCa. These chemo-sEVs contain a specific cargo profile that promotes the transfer of chemoresistance to sensitive tumoral cells (Bandari et al., 2020; Safaei et al., 2005; Tian et al., 2022; Yáñez-Mó et al., 2015). However, the underlying mechanisms implicated in facilitating biogenesis and secretion of chemo-sEVs in OvCa are poorly understood.

sEVs encompass a variety of vesicle types, including exosomes, that are formed intracellularly as intraluminal vesicles (ILVs) by the inward budding of the endosomal limiting membrane during the process of multivesicular endosomes (MVEs) maturation, organelles that fused with the PM causing the secretion of sEVs to the extracellular space (van Niel et al., 2018). The biogenesis of ILVs is mediated by endosomal sorting complex required for transport (ESCRT) machinery and mechanisms supported by specific lipids and scaffold proteins (Ceramide-Sphingomyelin, Tetraspanins, and Syndecan–Syntenin–ALIX dependent pathways) (Andreu & Yáñez-Mó, 2014; Baietti et al., 2012; Verderio et al., 2018; Wollert & Hurley, 2010). The transport, tethering, and fusion of MVEs with the PM are regulated by the RAB family small GTPases, molecules that cycle between GTP-bound active and GDP-bound inactive states (Cavieres et al., 2020; Kelly et al., 2012). RAB27A is one of the most studied small GTPases in this process and controls the trafficking and fusion of MVEs with the PM and the release of sEVs under basal and inducible conditions (Dorayappan et al., 2018; Ostrowski et al., 2010). Interestingly, RAB27A has been demonstrated to promote chemoresistance in different types of cancer models by a mechanism related with its role in exosomal secretion (Hertzman Johansson et al., 2013; J. Li et al., 2017; X. Li et al., 2017; J. Liu et al., 2017). However, whether CDDP-chemoresistance in OvCa is dependent on RAB27A function, remains unexplored.

MVEs are also key organelles in the endo-lysosomal degradative pathway. Here, MVEs fused with the lysosomes to degrade the cargo that gets incorporated into ILVs (Huotari & Helenius, 2011; Klumperman & Raposo, 2014), a pathway governed by RAB7 (Bucci et al., 2000; Vanlandingham & Ceresa, 2009), a small GTPase that contributes to the maintenance and biogenesis of lysosomes (Bae et al., 2019; Huotari & Helenius, 2011). The mechanisms that control the trafficking of MVEs to either the PM or in route to lysosomes are only partially understood (Eitan et al., 2016). It is well established that inhibiting the trafficking of MVEs in one direction can stimulate the alternative pathway as a compensatory response (Adams et al., 2021; Eitan et al., 2016; Guix et al., 2021; D. Huang et al., 2022; Miranda et al., 2018; van de Vlekkert et al., 2019; Villarroya-Beltri et al., 2016; J. Zhang et al., 2021), In the context of controlling chemo-sEVs secretion in OvCa cells, this means that targeting one MVE trafficking pathway (either secretory or lysosomal degradation) can have an impact on the balance between the two and potentially influence the secretion of chemo-sEVs. Particularly, it is proposed that this regulation is dependent on lysosome function (Izco et al., 2022). In this way, the contribution of lysosomes on sEVs secretion in cisplatin resistance in OvCa cells has been poorly explored.

Here, we studied the role of MVEs, ILVs, and sEVs in the chemoresistance of OvCa characterizing a cellular model sensitive (A2780) and chemo-resistant to CDDP (A2780cis). In comparison to A2780, A2780cis show an increased number of ILVs and higher levels of RAB27A, correlating with a higher sEVs secretion and transfer of chemoresistance capacity in response to CDDP. Conversely, we found A2780cis cells have a poor lysosomal function and low protein levels of RAB7. Collectively, these findings suggest that the high number of MVEs and ILVs found in A2780cis conduct sEVs secretion in a scenario of lysosomal dysfunction. Importantly, silencing of RAB27A converts A2780cis cells into an A2780-like phenotype promoting lysosomal function and restoring levels of RAB7. To mimic the effects of RAB27A silencing, the cells were treated with rapamycin, which promotes lysosomal biogenesis. The results showed that co-treating CDDP with rapamycin blocks the secretion of chemo-sEVs. In conclusion, we propose that the function of chemo- sEVs in OvCa is determined by an imbalance between MVE trafficking pathways, where the levels of RAB27A and RAB7, as well as the lysosomal status, controlled by mTORC1, play critical roles.

## Materials and Methods

### Antibodies

The following monoclonal antibodies were used: anti-ALIX (cat# sc-53540), anti- RAB7 (cat# sc-376362), anti-RAB11A (cat# sc-166912), anti-TSG101 (cat# sc- 7964) and anti-β-actin (cat# sc-47778) from Santa Cruz Biotechnology, Dallas, TX, USA; anti-CD63 (cat# ab8219), anti-GRP94 (cat# ab90458) and anti-RAB22A (cat# ab137093) from Abcam, Cambridge, UK; anti-LAMP1 clone H4A3 (cat# DSHB- H4A3) from Developmental Studies Hybridoma Bank, Iowa City, IA, USA; anti- TSG101 (cat# 612696) from BD Bioscience, Becton, NJ, USA. The following polyclonal antibodies: anti-Cathepsin D (cat# AF1014) from R&D Systems, Minneapolis, MN, USA; anti-CD63 (cat# AM11837) from Ango, San Ramon, CA, USA; anti-HGS (HRS) (cat# ab15539) from Abcam; anti-LAMP2A clone AMC2 (cat# 51-2200) from ThermoFisher Scientific, Waltham, MA, USA; anti-RAB35 (cat# 9690) from Cell Signaling Technology, Danvers, MA, USA; anti-RAB27A (cat# 168013) from Synaptic Systems, Göttingen, Germany. Horseradish peroxidase-conjugated secondary antibodies were purchased from Jackson ImmunoResearch Laboratories (West Grove, PA, USA), and Alexa fluorophore-conjugated secondary antibodies were purchased from Invitrogen^TM^, ThermoFisher Scientific, Carlsbad, CA, USA.

### Plasmids and oligos

The shRNA for RAB27A was cloned in pLKO.1 and provided courtesy of Matias Ostrowski Ph.D., Buenos Aires, Argentina (49). As a control we used a shRNA against Luciferase (cat# SHC007, Sigma-Aldrich, Saint Louis, MO, USA). The following sequences were used as a shRNA target shRNA-Luciferase: 5- ’CGCTGAGTACTTCGAAATGTC-3’, and shRNA-RAB27A: 5’-CGGATCAGTTAAGTGAAGAAA-3’. The packaging plasmids pCMV-*VSV-g* (cat# 8454) and pS-*PAX2* (cat# 12260) were obtained from Addgene (Watertown, MA, USA).

### Cell culture

HEK 293T cell line (Lenti-X™ 293T, Takara Bio, San Jose, CA, USA) was cultured in DMEM High Glucose (DMEM-HG cat# SH30243.02; Cytiva, Marlborough, MA, USA), supplemented with 10% heat-inactivated FBS (Sartorius, Göttingen, Niedersachsen, Germany), penicillin 100 U/mL, and streptomycin 100 mg/mL (Gibco^TM^, ThermoFisher Scientific, Waltham, MA, USA). A2780 (CDDP-sensitive) and A2780cis (CDDP-resistant) cell lines (Sigma-Aldrich) were cultured in RPMI 1640 (cat#SH30255.02, Cytiva, Marlborough, MA, USA), supplemented with 10% heat-inactivated fetal bovine serum, penicillin 100 U/mL and streptomycin 100 mg/mL. Every two weeks in culture or every three passages, A2780cis cells were treated with 1 μM CDDP (cat# 479306, Sigma-Aldrich,) for 72 h before seeding. The three cell lines were cultured at 37°C, in a humid environment, and 5% CO2 until the required cell confluence was obtained. Each 2-3 days, cells were trypsinized using Trypsin (0.05% Trypsin, 1 mM EDTA; Gibco^TM^, ThermoFisher Scientific) at 37°C. Alternatively, they were frozen in 1.5 mL of freezing medium (10 % v/v DMSO, 90 % v/v FBS) and stored in cryotubes at -80°C.

### Bacterial transformation and plasmid DNA isolation

One Shot^TM^ Stbl3^TM^ *E.coli* (Invitrogen^TM^, ThermoFisher Scientific, Carlsbad, CA, USA) was transformed with pLKO.1 vector, according to the manufacturer’s protocol. The transformed bacteria were grown to plasmid DNA isolation with NucleoBond Xtra Midi kit (Machery-Nagel, Allentown, PA, USA) according to the manufacturer’s protocol. The plasmid DNA isolated was quantified using the spectrophotometer EPOCH2 microplate reader (BioTek, Winooski, VT, USA).

### Generation of stable Knockdown cell lines with shRNA lentiviral particles

Lentiviral particles were generated by co-transfection of HEK293T cells with pLKO.1- *shRNA*, pCMV-*VSV-g*, and pS-*PAX2* vectors, harvested and concentrated as previously described (Varas-Godoy et al., 2018). The silencing of RAB27A was generated by transduction with shRNA-containing lentiviral particles. Cells were transduced with a shRNA against the luciferase to use as a control. For transduction, cells were infected and after 48 h selected and expanded with 6 μg/mL puromycin (Sigma-Aldrich).

### Western blot analysis

The preparation of cell lysate protein extracts, SDS-PAGE gels, and Western blot analysis was performed as previously described (Vargas et al., 2022). For the characterization of small extracellular vesicles, 10 µg of sEVs separated by SDS-PAGE gels were transferred to polyvinylidene difluoride membrane (PVDF; ThermoFisher Scientific) for 1 h at 100 V. Further, the membrane was analyzed using protocols previously described (Vargas et al., 2022; Vera et al., 2019).

### Transmission electron microscopy

Cells were grown on 60 mm cell culture plates and processed for transmission electron microscopy (TEM) analysis as previously described(Vargas et al., 2022). Isolated sEVs were fixed with paraformaldehyde (PFA), deposited in a TEM Formvar carbon grid, and contrasted with uranyl acetate as previously described (Acuña et al., 2020; Vera et al., 2019). Samples were analyzed using a Philips Tecnai 12 microscope (Eindhoven, The Netherlands) at 80 kV located in the Advanced Microscopy Facility of the Faculty of Biological Sciences at the Pontificia Universidad Católica de Chile.

### Indirect immunofluorescence and fluorometric lysosomal assays with fluorescent probes

Cells were seeded on glass coverslips pre-treated with L-poly-Lysine (cat# 4707 Sigma-Aldrich). After, cells were washed with PBS-Ca^2+^/Mg^2+^, fixed, permeabilized, and stained using our protocols previously published (Bustamante et al., 2020; Cavieres et al., 2020). The measurement of lysosomal pH and Cathepsin B activity was evaluated with LysoTracker^TM^ Red DND-99 probe (Invitrogen^TM^, ThermoFisher Scientific) and the Magic Red**®** kit (Immunochemistry Technologies, Davis, CA, USA), respectively, as previously reported (Vargas et al., 2022).

### Confocal fluorescence microscopy

Fluorescence microscopy images were acquired using a TCS SP8 laser-scanning confocal microscope (Leica Microsystems, Wetzlar, Germany) equipped with a 63x oil immersion objective (1.4 NA) running the Leica Application Suite LAS X software. The measurements were executed using ICY software (Quantitative Image Analysis Unit, Institut Pasteur, http://icy.bioimageanalysis.org/). A pipeline was created to completely automate image analysis by using the following sequential plugins: active contours (cell segmentation), hk-means (threshold detection), and wavelet spot detector (spot detection). For quantification of Manders coefficient colocalization, we used FIJI software version 2.1.0 (http://imagej.net/ software/fiji/) (Schindelin et al., 2012) plus the Just Another Colocalization Plugin (JACoP, 2.1.1 version) (Bolte & Cordelières, 2006), adjusting the threshold of the images respect to control.

### Small extracellular vesicles isolation

For sEVs isolation cells were cultured for 72 h in RPMI 1640 medium supplemented with 1% pen-strep and 5% exosome-depleted FBS (Gibco^TM^, ThermoFisher Scientific). sEV were isolated from the conditioned medium (CM) by differential ultracentrifugation (dUC), as previously described (Vera et al., 2019) using the SureSpin™ 630 Swinging Bucket Rotor (ThermoFisher Scientific).

### Small extracellular vesicles characterization and analysis

sEVs were characterized by Western blot, TEM, and nanoparticles tracking analysis (NTA) as previously (Vera et al., 2019). NTA analysis was performed using NanoSight NS300 equipment (Malvern Instruments, Malvern, United Kingdom, Universidad de los Andes). The measurement was performed using the 532-nm laser and 565-nm long pass filter, with a camera level at 9 and a detection threshold of 3. Small EVs were diluted in 1X PBS before the analysis.

### Chemoresistance transference and cell viability assays

For these assays we used a ratio of 50,000 sEVs per cell. After 16 h A2780 sensitive cells were treated with 3 µM CDDP for 48 h. Cell viability of A2780 cells after CDDP treatment was evaluated by the Live/Dead Cell Stain Kit (Invitrogen^TM^, ThermoFisher Scientific) by flow cytometry (BD FACS CantoII, Becton, NJ, USA) and using 0.4% trypan blue (cat# 15250-061, Gibco^TM^, ThermoFisher Scientific).

### Mass Spectrometry analysis of small extracellular vesicles

sEVs samples were compared by SWATH mass spectrometry (MS) to identify differentially expressed proteins. Briefly, 50 µg of sEVs proteins were mixed with RIPA buffer (pH 7.4), sonicated for 15 min, and centrifuged at 21,000 x g for 2 min at 4 °C. The proteins extracted from the sEVs were clean-up and concentrated using a 10 kDa Omega filtration centrifuge tubes (PALL, Nanosep), reduced, alkylated, and digested using the filter-aided sample preparation method (Wiśniewski et al., 2009). Post-digestion, 100 μL of 0.1 % formic acid (FA) was added to the digested sample and then loaded to a 10-kDa-size exclusion membrane (PALL, Nanosep), and centrifuged at 15,000 x g for 5 min to filter out peptides. The flow-through was reconstituted in an equal volume of 0.1 % trifluoroacetic acid (TFA) in preparation for desalting. The samples were desalted using SOLAμ HRP solid phase extraction (SPE) 96-well plate (ThermoFisher Scientific) according to the manufacturer’s instructions, and the desalted peptides were dried and reconstituted with 50 μL of 0.1 % FA in water.

Peptides were separated by liquid chromatography in tandem with MS (LC–MS) on an AB SCIEX 5600 Triple TOF spectrometer (ABSCIEX) coupled to a Nano Ultra 1D+ HPLC system (Eksigent). Each sample was loaded onto a reversed phase trap column (CHROMXP C18CL 5 μm, 10 x 0.3 mm; Eksigent) and the column wash was performed for 15 min (3 μL/min) followed by peptide separation on reversed phase analytical column (CHROMXP C18CL 3 μm, 120A0, 150 x 0.075 mm, Eksigent). The peptides were separated using a linear gradient with buffer A (H2O/0.1% FA), and buffer B (Acetonitrile (ACN)/0.1% FA) (75 min from 10% to 95% mobile phase B at 250 nL/min). Peptides were introduced into MS via a nanospray ion source (Thermo Fisher). The resulting peptides were processed in an information dependent acquisition (IDA) on an AB Sciex TripleTOF 5600 System using an isolation width of 26 Da (25 Da of optimal ion transmission efficiency and 1 Da for the window overlap), a set of 32 overlapping windows was constructed covering the mass range 100– 2000 m/z.

All mass spectra were analyzed using the Paragon algorithm on ProteinPilot™ Software (V4.5 beta, AB Sciex,) against a *Homo sapiens* Uniprot database. The change in the relative abundance of proteins in sEVs was established by comparing the extracted-ion peak intensities of two technical replicates of each sample using the SWATH Acquisition Microapp (version 2.0) within PeakView (version 2.2) (Alharbi et al., 2021). False discovery rate (FDR) analysis selected in the processing method which sets the detected protein threshold to 0.05 (10%). Venn diagram illustrating the shared proteins identified in the different sEVs groups was performed using the BioInfoRX Venn Diagram Plotter (https://bioinforx.com/apps/venn.php). Data was subjected to Gene Set Enrichment Analysis (GSEA) and Gene ontology analysis using the Webgestalt (http://www.webgestalt.org/), and the Protein-Protein Interaction Networks Functional Enrichment Analysis were evaluated using STRING (https://string-db.org/). The mass spectrometry work was performed at the University of Queensland Center for Clinical Research (UQCCR), Brisbane, Australia.

### Quantifications and Statistical Analysis

Densitometric quantification of images obtained by Western blot was estimated using FIJI software (Schindelin et al., 2012), version 2.1.0 (http://imagej.net/ software/fiji/). For each condition, protein bands were quantified from at least three independent experiments. Data analysis from densitometric quantifications, real-time qPCR, electron microscopy transmission, fluorescence confocal microscopy, and SRB-LC50 were performed using Microsoft Excel 2022 (Microsoft Corporation) and Prism 9.0 (GraphPad Software) for macOS Big Sur to generate corresponding charts and perform statistical analyses. Results are represented in graphs depicting the mean ± standard error of the mean (SEM). Statistical significance was determined by parametric one-tailed t-test and non-parametric one-tailed Mann-Whitney test, as indicated in each figure. P-values of *p<0.05, **p<0.01, and ***p<0.001 were regarded as statistically significant and are indicated in the figures. For mass spectrometry unpaired Student’s t-tests were performed with *p* < 0.05 considered statistically significant.

## Results

### A2780cis cells have increased protein machinery for ILVs biogenesis and exosome secretion

Secretion of sEVs<200 nm, has been suggested to mediate CDDP-chemoresistance in OvCa (Guerra et al., 2019; Safaei et al., 2005). However, whether this phenomenon is related to changes in MVEs and ILVs biogenesis has not been studied. To investigate whether this chemoresistance phenotype could be related to the amount of MVEs and ILVs biogenesis in A2780 CDDP-resistant (A2780cis), we analyzed CD63 positive punctated structures, a marker of MVEs compartments (Mathieu et al., 2021), in comparison to A2780 CDDP-sensitive (A2780) OvCa cells (Fig. 1A). We found by immunofluorescence confocal analysis that A2780cis cells have a higher number of CD63 positive punctated structures per cell than A2780 cells (Fig. 1A and 1B). In addition, CD63 positive punctated structures show an increased area in A2780cis cells compared to A2780 cells (Fig. 1A and 1C). This initial finding prompted us to investigate whether A2780cis could have a higher capacity in MVEs and ILVs biogenesis (Peng et al., 2021). Specifically, we studied MVEs and ILVs structures by transmission electron microscopy (TEM) in both cell lines (Fig 1D). We found in A2780cis cells that MVEs organelles showed an increased area in comparison to A2780 cells (Fig. 1E). Moreover, we observed a higher number of ILVs per MVEs in A2780cis compared to A2780 cells (Fig. 1F). These results strongly suggest that A2780cis cells have an increased capacity to form MVEs and ILVs from *novo* synthesis. To evaluate this possibility, we measured the levels of ESCRT family members involved in ILVs biogenesis including ALIX, HRS, and TSG101 (Falguières et al., 2008; Wollert & Hurley, 2010). Additionally, we assessed levels of LAMP2A, a transmembrane protein recently linked to the biogenesis of ILVs and secretion of sEVs (Ferreira et al., 2022). Our western blot analysis revealed that all the proteins evaluated had significantly higher levels in A2780cis cells, compared to A2780 cells (Fig. 1G and 1H). This confirms that CDDP- resistant cells have enhanced protein machinery necessary for ILVs biogenesis. Given the increase in the size of the MVEs and the number of ILVs, which may imply an increased capacity for sEVs secretion, we next examined the levels of key RAB GTPases known to play critical roles in the secretion of sEVs (Jin et al., 2021; Messenger et al., 2018; Ostrowski et al., 2010; T. Wang et al., 2014; Wei et al., 2021). Our results showed that among the RABs evaluated (RAB11A, RAB22A, RAB27A, RAB35), only the levels of RAB27A levels increased significantly in A2780cis compared to A2780 cells (Fig. 1I and 1J). Since RAB27A has been linked to the secretion of sEVs (Ostrowski et al., 2010), these findings suggest that the CDDP-resistant phenotype in OvCa could be associated with an increased ability of these tumoral cells to increase the biogenesis of MVEs and ILVs, leading to a more secretory state and increased secretion of sEVs.

**Figure 1.**
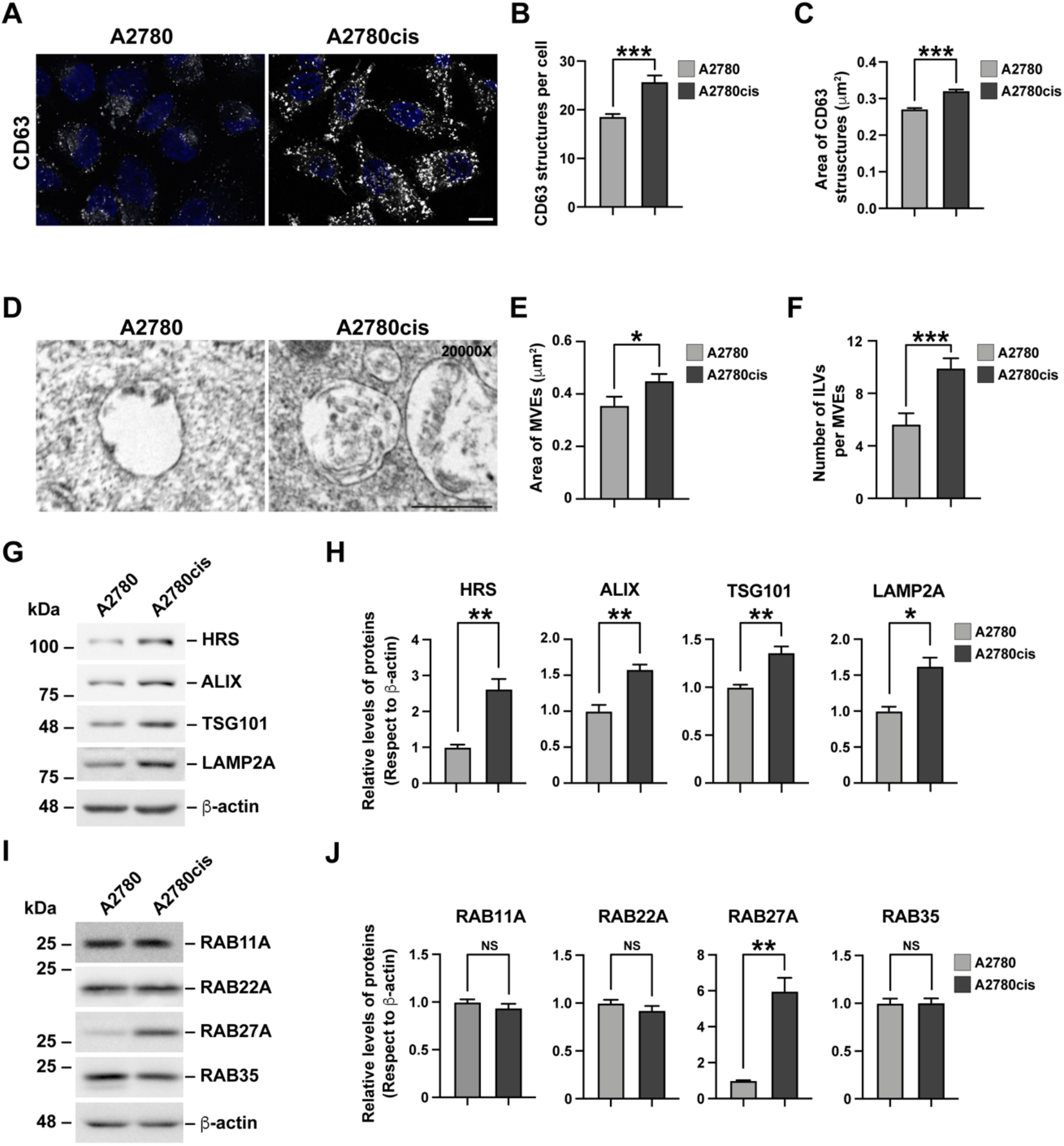
A2780cis CDDP-Resistant cells have more capacity to produce and secrete exosomes. **(A)** Representative confocal microscopy images of A2780 and A2780cis PFA-fixed cells immunofluorescent stained with anti-CD63. Scale Bar 10 μm. **(B)** Analysis of the number of structures and **(C)** area of structures of CD63 per cell from images as those shown in (A). Bars indicated the mean with SEM; ***P<0.001; one-tiled unpaired parametric t-Test (n>50 cells). **(D)** Representative TEM micrograph shows MVEs of A2780 and A2780cis cells. Scale Bar 0.5 μm. **(E)** Semiquantitative analysis of the area of MVEs and **(F)** ILVs per MVEs of A2780 and A2780cis cells, from images as those shown in (D). Bars indicated the mean with SEM; *P<0.05, ***P<0.001; one-tiled, unpaired, parametric t-Test (n>50 cells). **(G)** Analysis of detergent-soluble protein extracts obtained from A2780 and A2780cis cells by western blot with anti-HRS, anti-ALIX, anti-TSG101, anti-LAMP2A and anti- β-actin. The image is representative of three independent experiments. **(H)** Densitometric quantification of the signal of HRS, ALIX, TSG101, and LAMP2A from images as those shown in (G). The signal was normalized with β-actin signal. Bars indicated the mean with SEM; *P<0.05, **P<0.01, one-tiled, paired, non-parametric Mann-Whitney test (n = 3). **(I)** Analysis of detergent-soluble protein extracts obtained from A2780 and A2780cis cells by western blot with anti-RAB11A, anti-RAB22A, anti-RAB27A, anti-RAB35, and anti-β-actin. The image is representative of three independent experiments. **(J)** Densitometric quantification of the signal of RAB11A, RAB22A, RAB27A, and RAB35 from images as those shown in (I). The signal was normalized with β-actin signal. Bars indicated the mean with SEM; **P<0.01, NS: Not significant; one-tiled, paired non-parametric Mann-Whitney test (n = 3).

### sEVs secreted from A2780cis cells in response to CDDP can transfer cisplatin chemoresistance to recipient A2780 cells

Because A2780cis cells showed a higher molecular capacity for MVEs and ILVs biogenesis, we investigated if this feature could correlate with the secretion of sEVs which include exosomes. For this, we isolated sEVs by differential centrifugation from conditioned media of A2780 and A2780cis cells cultured for 72 h (Vera et al., 2019). Additionally, we isolated sEVs from A2780cis cells treated with 1 μM CDDP for 72 h. A2780 cells in the presence of CDDP were not evaluated considering most cells died after this treatment. In addition, it was previously demonstrated that this drug for a short time of incubation does not induce the secretion of sEVs in A2780 cells (Samuel et al., 2018). First, we characterized if the isolated sEVs fractions were enriched on selected exosome markers according to current recommendations (MISEV 2018) (Théry et al., 2018). As expected, we found the sEVs fractions isolated from the three different conditioned media were enriched in endosomal proteins ALIX, TSG101, and CD63, respecting the whole total cell lysate (Fig. 2A). As an internal sEVs purity control, we measured the levels of the endoplasmic reticulum (ER) luminal protein GRP94, an abundant protein known to be absent in sEVs fractions (Dozio & Sanchez, 2017). Importantly, our analysis showed GRP94 was detected only in the cell lysate fractions (Fig. 2A). Next, to analyze the distribution size of the isolated sEVs fractions in each condition, we performed Nanoparticle Tracking Analysis (NTA) analysis. We found all isolated sEVs were enriched in particles between 120 to 200 nm in diameter (Fig. 2B). Moreover, the microphotographs show that all sEVs fractions have the typical cup shape morphology (Fig. 2C). Collectively, the protocol selected allows the isolation of pure sEVs enriched in exosome markers (Théry et al., 2018), hereafter indicated for simplicity sEVs. Next, we measured the number of sEVs in each condition by NTA analysis. Unexpectedly, we found that A2780 and A2780cis secreted a similar number of sEVs per cell (Fig. 2D). However, treatment of A2780cis with CDDP caused a significant increase in the number of secreted sEVs per cell, compared with untreated cells (Fig 2D). In this regard, it is known that cancer cells secrete sEVs in response to chemotherapeutic agents such as CDDP named as chemo- sEVs (Ab Razak et al., 2019; Bandari et al., 2018), which can mediate horizontal transference of drug resistance to drug-sensitive cells (Xavier et al., 2022; Yáñez-Mó et al., 2015).

**Figure 2.**
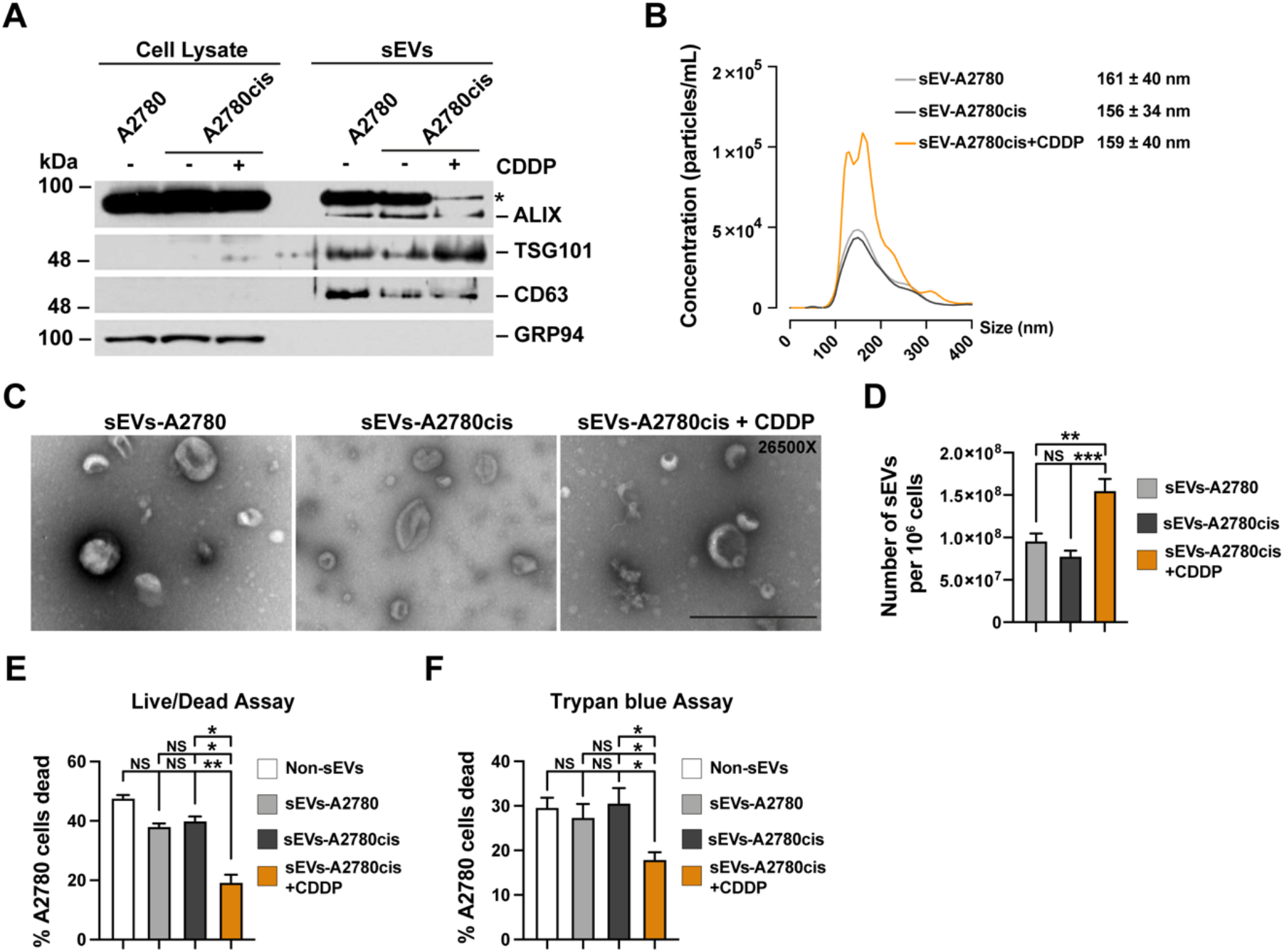
CDDP promotes the secretion of small EVs in A2780cis CDDP-resistant OvCa cells, with chemoresistance transference capacity. **(A)** Analysis of detergent- soluble protein extracts by western blot from cell lysates and small EVs (sEVs) obtained by differential ultracentrifugation (dUC) from 72 h conditioned medium (CM) of A2780 (sEVs-A2780), A2780cis (sEVs-A2780cis) and A2780cis cells treated with 1 μM CDDP during 72 hrs (sEVs-A2780cis+CDDP). To western blot, we used anti- ALIX, anti-TSG101, anti-CD63, and anti-GRP94. * Indicate unspecified band. **(B)** Nano-tracking analysis (NTA) of sEVs-A2780, sEVs-A2780cis, and sEVs- A2780cis+CDDP obtained by dUC. The quantifications represent the distribution size of the sEVs calculated by the mean of concentration of the sEVs of each size (n = 5). **(C)** Representative images of TEM micrograph of sEVs-A2780, sEVs- A2780cis, and sEVs-A2780cis+CDDP, obtained by dUC. ScaleBar 0.5 μm. **(D)** Analysis of the number of sEVs-A2780, sEVs-A2780cis, and sEVs-A2780cis+CDDP per 106 cells obtained by dUC. Bars indicated the mean with SEM; **P<0.01, ***P<0.001, NS: Not significant; one-tiled, unpaired, non-parametric Man-Whitney test (n = 5). **(E-F)** Live/Dead cell stain analysis by (E) flow cytometry and (F) trypan blue analysis of A2780 cells treated with 3 µM CDDP for 48 hours after 16 hours stimulation with PBS (Non-sEVs) or with sEVs-A2780, sEVs-A2780cis, and sEVs- A2780cis+CDDP obtained by dUC. The analysis represents the percentage of A2780 cells dead for each condition. Bars indicated the mean with SEM; *P<0.05, **P<0.01, NS: Not significant; one-tiled, paired, non-parametric Mann-Whitney test (n = 3).

This feature prompted us to evaluate the potential of these sEVs secreted in response to CDDP, in their capacity to transfer CDDP resistance to chemo-sensitive cells. To do this, we treated A2780cis cells with 1 μM CDDP for 72 h to generate chemo-sEVs and then tested their ability to confer chemo-resistance when incubated with A2780 cells at a ratio of 50,000 chemo-sEVs per cell for 16 h. After, cells were treated with 3 µM CDDP for 48 h, a concentration that caused approximately 50% mortality after the treatment. Subsequently, the viability of the cells was assessed using both live/dead cell staining analysis by flow cytometry and trypan blue analysis. As controls, A2780 cells were incubated with 50,000 sEVs per cell, which were isolated from either A2780 cells or A2780cis, but without the addition of CDDP. Our results reveal that the chemo-sEVs fraction was effective in significantly reducing CDDP-induced death in A2780 cells compared to the other sEVs fractions, as shown by live/dead staining analysis (Fig. 2E) and trypan blue analysis (Fig. 2F). Therefore, our findings demonstrated the transfer of chemo-sEVs resistance from A2780cis cells by the sEVs to A2780 chemo-sensitive cells, as indicated by the reduction of CDDP-induced cell death.

### sEVs secreted from A2780cis cells in response to CDDP are enriched in proteins involved in chemoresistance

To get insights about which key components of chemo-sEVs could be implicated in the transfer of CDDP chemo-resistance in A2780 chemo-sensitive cells, we performed SWATH mass spectrometry (MS) to identify specific proteins present in those sEVs isolated from A2780cis in response to CDDP (Fig. 3). A total of 330, 578, and 479 proteins were identified in the sEVs fractions isolated from A2780, A2780cis, and A2780cis+CDDP, containing 37, 167, and 84 proteins exclusive in each group respectively (Fig. 3A and Supplementary Table S1). Gene ontology analysis revealed that the proteomic profile of the different sEVs are mainly related to pathways involved in cell communication, signal transduction, and metabolism (Fig. 3B). A quantitative proteomic analysis shows 65 proteins overrepresented in the chemo-sEVs in comparison with the sEVs derived from the A2780cis (Fig. 3C and Supplementary Table S2). A protein-protein interaction network and functional enrichment analysis of the total proteins found in the chemo-sEVs reveal a network of proteins associated with DNA Double-strand break repair (CHEK2, PRKDC, HIST1H4F, HIST1H2BM, XRCC6, UBA52), proteins of the proteasome complex which are involved in stabilization and regulation of p53, RUNX2 and PTEN (PSMC2, PSMA7, PSMA5, PSMD12, PSMC3, PSMC5, PSMD11, PSMD14, PSMC6, PSMD7, and PSME3), and proteins associated to the metabolism of RNA (mostly ribosomal proteins) (Fig. 3D). All these networks have been associated with the acquisition of chemoresistance, therefore this functional network analysis reveals possible mechanisms of how chemo-sEVs could transfer chemoresistance to sensitive cells as we previously described in this work (Fig. 2E and 2F).

**Figure 3.**
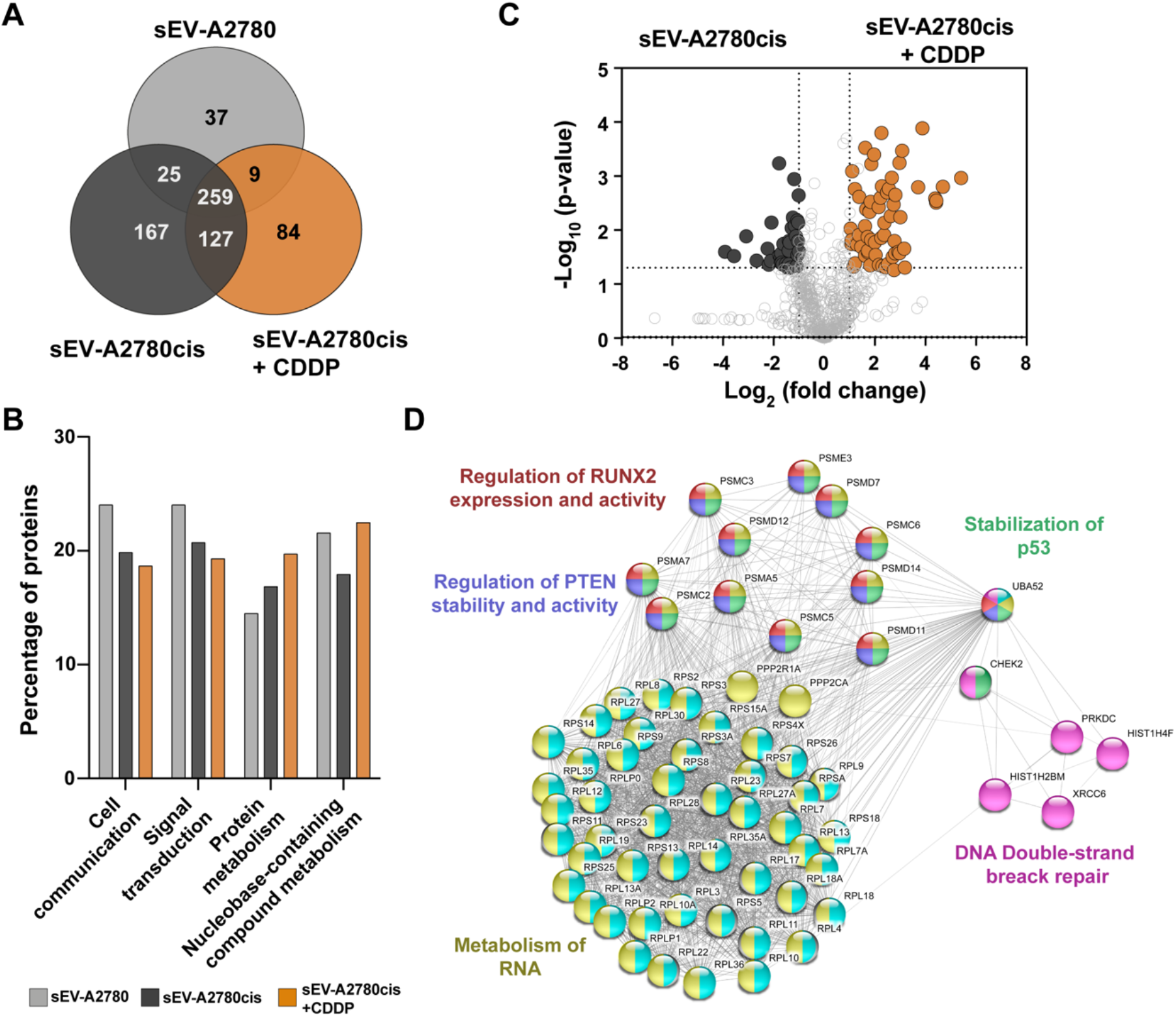
Small EVs protein cargo of A2780cis cells is altered after treatment with CDDP and is enriched with proteins associated with chemoresistance. **(A)** Venn diagram of proteins identified in sEVs obtained by dUC from 72 h CM of A2780 (sEVs-A2780), A2780cis (sEVs-A2780cis) and A2780cis cells treated with 1 μM CDDP during 72 hrs (sEVs-A2780cis+CDDP). **(B)** Gene ontology (GO) enrichment analysis of proteins found in sEVs-A2780, sEVs-A2780cis, and sEVs-A2780cis+CDDP obtained by dUC. **(C)** Volcano plot of proteins differentially expressed between sEVs-A2780cis, and sEVs-A2780cis+CDDP. **(D)** Protein-Protein Interaction Networks Functional Enrichment Analysis of total proteins differentially expressed in sEVs-A2780cis+CDDP.

### A2780cis cells present a lysosomal alteration phenotype

Further, we investigated the underlying cellular mechanisms that could explain the increased machinery of A2780cis to form MVEs and ILVs, key mediators for the major secretion of chemo-sEVs. In this context, and because previous findings in C13 cells, other OvCa CDDP-resistant cellular model, suggested that a high number of MVEs could be related with disturbances at the level of lysosomes (Guerra et al., 2019; Safaei et al., 2005), acidic organelles functionally associated with MVEs compartments (Peng et al., 2021), we tested the status of lysosomes between A2780 and A2780cis cells. First, we evaluated by western blot the protein levels of the structural lysosomal membrane protein LAMP1. Interestingly, we observed a significant decrease in levels of LAMP1 in A2780cis cells compared to A2780 cells (Fig. 4A and 4B). Then, we analyzed by immunofluorescence the number of lysosomes, using as a marker LAMP1 positive punctated structures positive to Cathepsin D staining, an acidic luminal hydrolase known as a classical marker of lysosomes (Oberle et al., 2010; Pi et al., 2017). By immunofluorescence we observed that A2780cis showed a notorious decrease in the number of punctated structures positive to both, LAMP1 and Cathepsin D (Fig. 4C). Quantitative analysis of these images confirmed a significant decrease in the number of LAMP1 (Fig. 4D) and Cathepsin D (Fig.4E) structures. Moreover, we found a significant reduction in the average intensity per punctated structures in A2780cis, either positive to LAMP1 (Fig.4F) and Cathepsin D (Fig. 4G), compared to A2780 cells. Next, we investigated whether the reduction in LAMP1 and Cathepsin D structures were indicative of lysosomal function impairment. For this, we performed labeling of acidic compartments with the LysoTracker™-Red probe in live cells (Fig. 4H). We observed a significant reduction in the number of acidic structures in A2780cis compared to A2780 (Fig. 4I). As observed with LAMP1 and Cathepsin D, the measurement of the average intensity of lysotracker positive structures also showed a significant decrease in A2780cis compared to A2870 cells (Fig. 4J). Moreover, because an acidic environment is critical for lysosomal hydrolase activities, we tested whether the reduction in lysosomal acidity in A2780cis cells correlated with a reduction in the activity of the Cathepsin B, an abundant lysosomal hydrolase (Oberle et al., 2010). For this, lysosomes were labeled with the Magic Red**®**, a probe that allows to quantify the activity of Cathepsin B in live cells (Kundu et al., 2018). As expected, the quantification of average intensity showed a significant reduction in Cathepsin B activity in A2780cis cells with respect to A2780 cells (Fig. 4K and 4L). Finally, we tested the RAB7 protein levels, a GTPase that mediates the trafficking of MVEs and their fusion with lysosomes, to the maintenance of this organelle (Bucci et al., 2000). Surprisingly, and contrary to our finding with RAB27A (Fig 1I and 1J), we found that levels of RAB7 were significantly decreased in A2780cis compared to A2780 cells (Fig. 4M and 4N). This decrease could indicate a lower fusion capacity of MVEs with lysosomes (Bucci et al., 2000).

**Figure 4.**
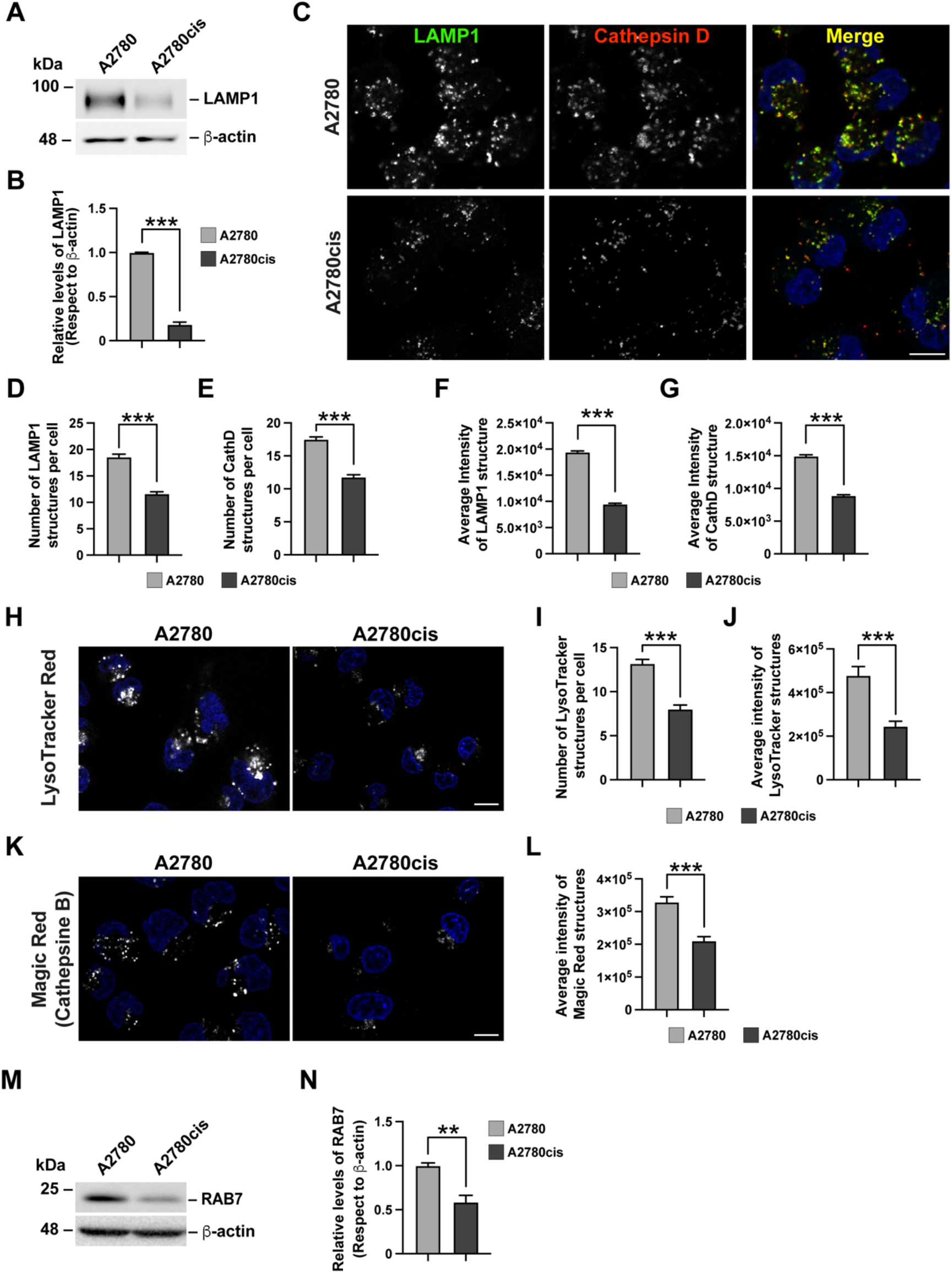
A2780cis CDDP-resistant OvCa cells have lysosomal alteration. **(A)** Analysis of detergent-soluble protein extracts obtained from A2780 and A2780cis cells by western blot with anti-LAMP1 and anti-β-actin. The image is representative of three independent experiments. **(B)** Densitometric quantification of the signal of LAMP1 from images as those shown in (A). The signal was normalized with β-actin signal. Bars indicated the mean with SEM; ***P<0.001; one-tiled, paired, non- parametric Mann-Whitney test (n = 3). **(C)** Confocal microscopy images of A2780 and A2780cis PFA-fixed cells immunofluorescent stained with anti-LAMP1 and anti- Cathepsin D. Scale Bar 10 μm. **(D-E)** Analysis of the number of structures LAMP1 (D) and Cathepsin D (CathD) (E) from images as those shown in (C). Bars indicated the mean with SEM; ***P<0.001; one-tiled, unpaired, parametric t-Test (n>50 cells). **(F-G)** Analysis of average intensity of LAMP1 (F) and CathD (G) structures from images as those shown in (C). Bars indicated the mean with SEM; ***P<0.001; one- tiled, unpaired, parametric t-Test (n>50 cells). **(H)** Confocal live-cell images of A2780 and A2780cis cells incubated with LysoTracker Red and HOECHST33342. Scale Bar 10 μm. **(I-J)** Analysis of number (I) and average intensity (J) of LysoTracker Red structures from images as those shown in (H). Bars indicated the mean with SEM; ***P<0.001; one-tiled, unpaired, parametric t-Test (n>50 cells). **(K)** Confocal live-cell images of A2780 and A2780cis cells incubated with Magic Red and HOECHST33342. Bar 10 μm. **(L)** Analysis of average intensity of Magic Red structures from images as those shown in (K). Bars indicated the mean SEM; ***P<0.001; one-tiled, unpaired, parametric t-Test (n>50 cells). **(M)** Analysis of detergent-soluble protein extracts obtained from A2780 and A2780cis cells by western blot with anti-RAB7 and anti-β-actin. The image is representative of three independent experiments. **(N)** Densitometric quantification of the signal of RAB7 from images as those shown in (M). The signal was normalized with β-actin signal. Bars indicated the mean with SEM; **P<0.01; one-tiled, paired, non-parametric Mann-Whitney test (n = 3).

Our findings showed that the CDDP chemo-resistant phenotype of A2780cis, which is characterized by high sEVs secretion, is probably by a decline in lysosomal degradative function and a decrease in the levels of RAB7 GTPase. This affects the trafficking and fusion of MVEs towards lysosomes leading to a shift in the trafficking route to the plasma membrane with a subsequent increase in sEVs secretion.

### Silencing of RAB27A reestablishes lysosomal function impairment in A2780cis cells

The increased secretion of sEVs in A2780cis cells may be a result of lysosomal dysfunction, as evidenced by the high number of MVEs and ILVs found in these cells which are specialized in the secretion of chemo-sEVs in response to CDDP. This phenotype is associated with a significant reduction in lysosomal activity, leading us to believe that MVEs may adopt this secretory status as a means of compensating for the impaired lysosomal function.

Moreover, the phenotype of increased sEVs secretion in response to lysosomal dysfunction in A2780cis cells is also associated with elevated levels of the RAB27A GTPase. Based on this correlation, we asked if silencing of RAB27A could be an effective approach to revert the lysosomal dysfunction. To test this possibility, we silenced RAB27A in A2780cis cells by stable expression of a specific shRNA (shRAB27A), a strategy that has been proven to specifically block the secretion of exosomes (Blanc & Vidal, 2018; Bobrie et al., 2012; H. Huang et al., 2021; Ostrowski et al., 2010; Salimu et al., 2017). As controls, we generated A2780 and A2780cis cells that stably expressed an shRNA against Luciferase (shLuc). By western blot we confirmed efficient silencing of RAB27A in A2780cis cells (Fig. 5A and 5B). Then, we evaluated if silencing of RAB27A could trigger some effect in lysosomes in A2780cis. First, we tested the levels of RAB7, a GTPase was found to be diminished in A2780cis. Surprisingly, silencing of RAB27A causes a significant increase in the levels of RAB7, reaching similar levels to those found in A2780 CDDP-sensitive cells (Fig. 5A and 5B). High levels of RAB7 are usually related to an increased number of lysosomes (Bucci et al., 2000). To test this possibility, we evaluate by Western blot the levels of LAMP1. Interestingly, we found that silencing of RAB27A enhances the levels of LAMP1 in A2780cis cells, again reaching similar levels to those found in A2780 CDDP sensitive cells (Fig. 5A and 5B). Next, we investigated if high levels of LAMP1 in A2780cis were related to an increase in the number of lysosomes. By immunofluorescence, we found silencing of RAB27A in A2780cis caused a strong increase in the number of lysosomes, measured by the increase in LAMP1 and Cathepsin D positive co-staining (Fig. 5C, lower panel compared to the middle panel). Importantly, we observed that silencing of RAB27A in A2780cis caused the recovery in lysosomal structures observing a pattern similar to A2780 cells (Fig. 5C, lower panel compared to the top panel). Quantification analysis confirmed a significant increase either in the number of LAMP1 (Fig. 5D) and Cathepsin D structures per cell (Fig. 5E) as well as in the average intensity of these LAMP1 (Fig. 5F) and Cathepsin D structures (Fig. 5G). Next, we evaluated if the enhancement in lysosomal structures was accompanied by an increase in the acidity and activity of these organelles. For this, we used the LysoTracker probe, which labels acidic lysosomal compartments. We observed that the reduced number and intensity of LysoTracker positive structures found in A2780cis cells, in comparison to A2780 sensitive cells, was significantly enhanced with the silencing of RAB27A to levels similar to those found in A2780 cells (Fig. S1A, S1B, S1C). In addition, we also found that silencing of RAB27A in A2780cis cells caused a significant increase in the average intensity labeling with Magic Red, observing similar levels to those found in A2780 cells (Fig. S1D and S1E). Finally, we evaluated levels of the tetraspanin CD63, a protein that is enriched in MVEs and ILVs compartments, characterized by the presence of a lysosome targeting motif in their sequence that allows its degradation by lysosomes (Kubo et al., 2017; Mathieu et al., 2021). With the rationale that silencing of RAB27A could promote more efficient trafficking of MVEs toward lysosomal degradation instead of secretion, we studied levels of CD63 by immunofluorescence. Surprisingly, we found that silencing of RAB27A in A2780cis cells significantly reduced the number and average intensity of CD63 positive structures, similar to those in A2780 CDDP-sensitive cells (Fig. 5F and 5G). These findings strongly suggest that silencing of RAB27A reverts the altered lysosomal phenotype of A2780cis cells, allowing MVEs to be trafficked towards lysosomal degradation instead of secretion, which is similar to A2780 CDDP-sensitive cells.

**Figure 5.**
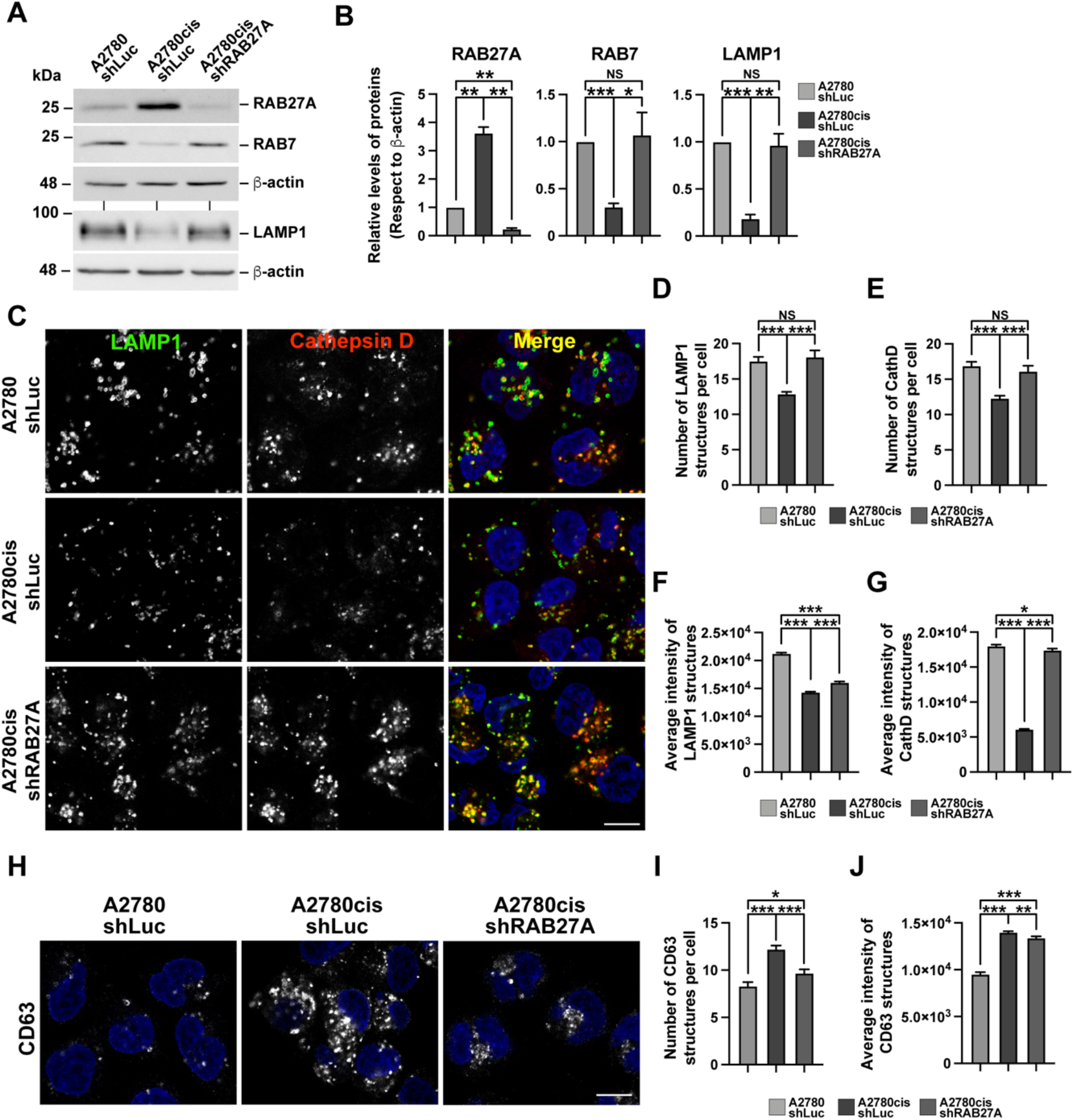
RAB27A silencing expression reverses lysosomal alteration phenotype in A2780cis CDDP-resistant OvCa cells. **(A)** Analysis of detergent-soluble protein extracts obtained from A2780 shLuc, A2780cis shLuc, and A2780cis shRAB27A stable cell lines by western blot with anti-RAB27A, anti-RAB7, anti-LAMP1, and β- actin. The image is representative of three independent experiments. **(B)** Densitometric quantification of the signal of RAB27A, RAB7, and LAMP1 from images as those shown in (A). The signal was normalized with β-actin signal. Bars indicated the mean with SEM; A2780 shLuc samples were normalized to 1 since the biological replicates were run in different western blot; *P<0.05, **P<0.01, ***P<0.001, NS: Not significant; one-tiled, paired, non-parametric Mann-Whitney test (n = 3). **(C)** Confocal microscopy images of A2780 shLuc, A2780cis shLuc, and A2780cis shRAB27A PFA-fixed cells immunofluorescent stained with anti-LAMP1 and anti-Cathepsin D. Scale Bar 10 μm. **(D-E)** Analysis of the number of structures of LAMP1 (D) and Cathepsin D (CathD) (E) from images as those shown in (C). Bars indicated the mean with SEM; ***P<0.001, NS: Not Significant; one-tiled unpaired t- Test (n>50 cells). **(F-G)** Analysis of average intensity of LAMP1 (F) and CathD (G) structures from images as those shown in (C). Bars indicated the mean with SEM; *P<0.05, ***P<0.001; one-tiled, unpaired, parametric t-Test (N>50 cells). The recovery of lysosomal acidity and activity is shown in Fig. S1. **(H)** Confocal microscopy images of A2780 shLuc, A2780cis shLuc, and A2780cis shRAB27A PFA-fixed cells immunofluorescent stained with anti-CD63. **(I-J)** Analysis of number (I) and average intensity (J) of CD63 per cell from images as those shown in (H). Bars indicated the mean with SEM; *P<0.05, **P<0.01, ***P<0.001; one-tiled, unpaired, parametric t-Test (n>50 cells).

### Reestablishment of lysosomal function in A2780cis cells reduces the secretion of chemo-sEVs and resistance transfer capacity

Since the silencing of RAB27A might promote the trafficking of MVEs toward degradative pathway, enhancing lysosomes function and impacting the secretion of chemo-sEVs (Fig. 5), prompted us to evaluate if another well-known lysosomal inducing condition would result in a similar phenotype. To address this, we tested the effect of rapamycin on A2780cis cells, an inhibitor of the mTORC1 kinase, which is known as a potent signaling pathway that induces lysosomal function (Palmieri et al., 2011; Settembre et al., 2012). First, we confirmed the inhibitory effect of rapamycin on mTORC1 activity by measuring the phosphorylation of the T389 residue in S6K, a substrate of mTORC1 (Rosner et al., 2012). The effect of rapamycin in A2780cis cells was tested in the absence or presence of CDDP treatment for 72 h. In agreement with a previous report, we observed that CDDP enhances mTORC1 activity demonstrated by an increase in S6K phosphorylation respect to untreated cells (Fig. 6A and 6B) (Gremke et al., 2020). Since hyperactivation of mTORC1 negatively impacts on lysosomal function (Palmieri et al., 2011; Settembre et al., 2012), it confirmed the hypothesis that the increased secretion of chemo-sEVs by CDDP treatment is due to a decline in lysosomal function. Rapamycin blocked the basal and CDDP-induced phosphorylation of S6K as expected (Fig. 6A and 6B). We then investigated if rapamycin could be able to recover the reduced number of lysosomes found in A2780cis cells. For this, A2780cis cells treated with rapamycin in the absence or presence of CDDP for 72 h were tested with antibodies against LAMP1 and Cathepsin D (CathD) by immunofluorescence. Rapamycin treatment alone or with CDDP caused an increase in the signal of LAMP1 and Cathepsin D in A2780cis cells as shown by immunofluorescence analysis (Fig. 6C). In contrast, no changes were observed in the presence of CDDP, observing a similar pattern that untreated cells (Fig. 6C). Quantification analysis confirmed a significant increase in the average intensity of structures positive to LAMP1 (Fig. 6D) and Cathepsin D (Fig 6E) in the presence of rapamycin. Following, we investigated the effect of rapamycin treatment on the secretion of sEVs by A2780cis in response to CDDP treatment (chemo-sEVs) for 72 h. First, we measured the distribution size of sEVs by NTA analysis in each condition.

**Figure 6.**
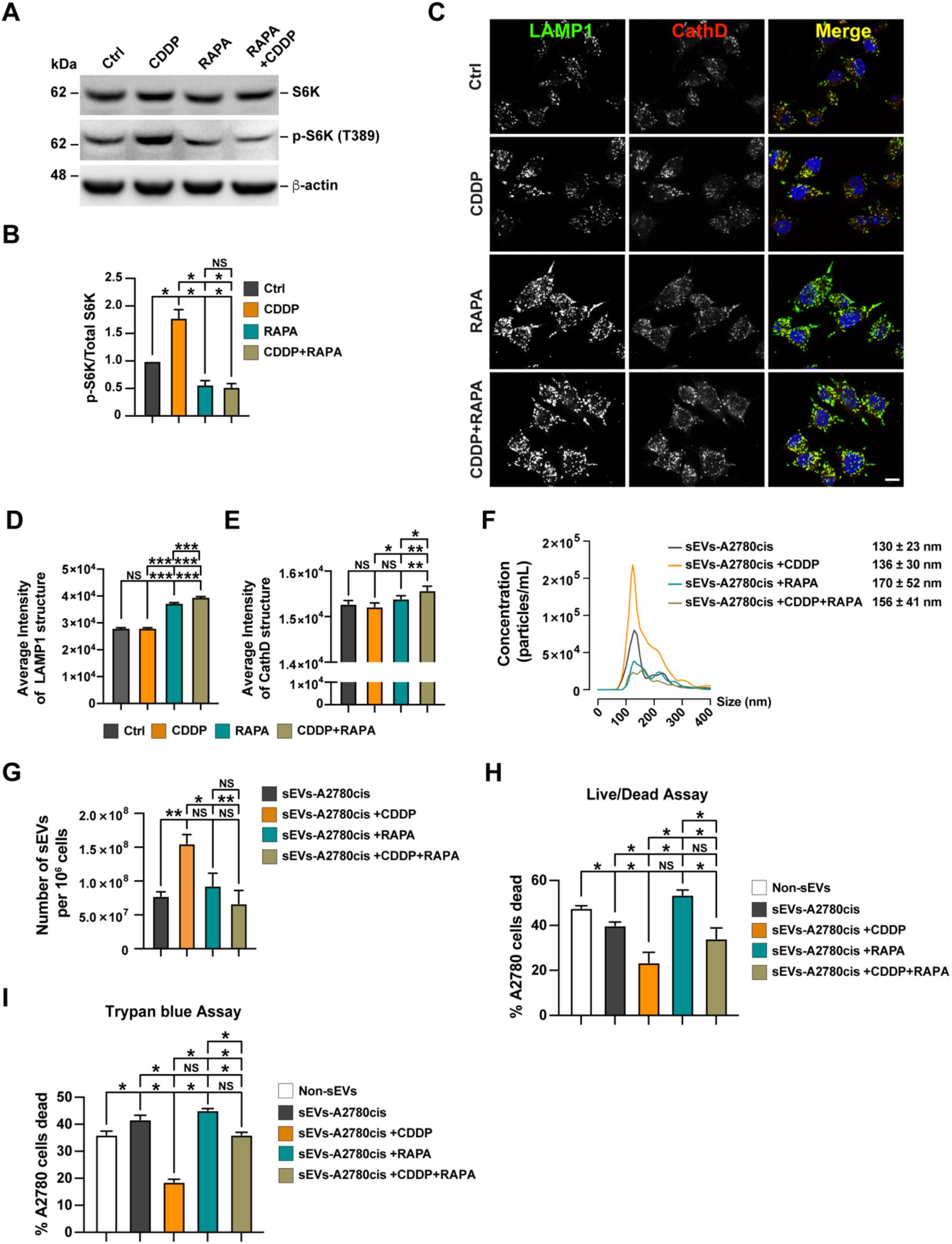
Induction of lysosomal biogenesis by rapamycin prevents CDDP effect on A2780cis CDDP-resistant OvCa cells in the secretion of sEVs and chemoresistance transference capacity. **(A)** Analysis of detergent-soluble protein extracts obtained from A2780cis control, treated with 1 μM CDDP, 100 nM rapamycin (RAPA), or 1 μM CDDP plus 100 nM RAPA during 72 hrs, by western blot with anti-S6K, anti-S6K (T389), and β-actin. **(B)** Densitometric quantification of the signal of p-S6K (T389) respect to total S6K. Bars indicated the mean with SEM; Control samples were normalized to 1 since the biological replicates were run in different western blot; *P<0.05, NS: Not significant; one-tiled, paired, non-parametric Mann-Whitney test (n = 3). **(C)** Confocal microscopy images of A2780cis control, treated with 1 μM CDDP, 100 nM RAPA or 1 μM CDDP plus 100 nM RAPA during 72 h. Scale Bar 10 μm. **(D-E)** Analysis of average intensity of structures of LAMP1 (D) and CathD per cell (E) from images as those shown in (C). Bars indicated the mean with SEM; *P<0.05, **P<0.01, ***P<0.001, NS: Not Significant; one-tiled, unpaired, parametric t-Test (n>50 cells). **(F)** NTA of sEVs obtained by dUC from 72 h CM of A2780cis (sEVs- A2780cis), A2780cis cells treated with 1 μM CDDP during 72 h (sEVs- A2780cis+CDDP), treated with 100 nM RAPA during 72 h (sEVs-A2780cis+RAPA), and treated with 1 μM CDDP plus 100nM RAPA during 72 h (sEVs- A2780cis+CDDP+RAPA). The quantifications represent the distribution size of the sEVs calculated by the mean concentration of the sEVs of each size (n = 5). **(G)** Analysis of the number of sEVs-A2780cis, sEVs-A2780cis+CDDP, sEVs- A2780cis+RAPA, and sEVs-A2780cis+CDD+RAPA per 10^6^ cells of obtained by dUC. Bars indicated the mean with SEM; *P<0.05, **P<0.01, NS: Not significant; one-tiled, unpaired, non-parametric Mann-Whitney test (n = 5). **(H-I)** Live/Dead cell stain analysis by flow cytometry (H) and trypan blue analysis (I) of A2780 cells treated with 3 µM CDDP for 48 hrs after 16 hrs of stimulation with PBS (Non-sEVs) or with sEVs-A2780cis, sEVs-A2780cis+CDDP, sEVs-A2780cis+RAPA, or sEVs- A2780cis+CDD+RAPA obtained by dUC. The analysis represents the percentage of A2780 cells dead for each condition. Bars indicated the mean with SEM; *P<0.05, NS: Not significant; one-tiled, paired, non-parametric Mann-Whitney test (n = 3).

This analysis showed no changes in this parameter, corresponding in all conditions to sEVs between 130 to 170 nm in diameter, confirming they correspond to sEVs (Fig. 6F). Moreover, we studied the number of sEVs in each condition. As previously, CDDP caused a significant increase in the secretion of sEVs (Fig. 6G). In contrast, rapamycin treatment did not affect the number of sEVs secreted by A2780cis cells, as shown in Fig. 6G. Importantly, rapamycin caused a potent inhibition in the number of sEVs secreted in response to CDDP (Fig. 6G). Finally, we studied if rapamycin could reduce the resistance transfer capacity of sEVs secreted in response to CDDP. As our previous assays, A2780 cells were incubated with 50,000 chemo-sEVs per cell for 16 h, which were isolated from the conditioned medium of A2780cis treated for 72 h in each condition. After incubation with chemo-sEVs, A2780 cells were treated with 3 µM CDDP for 48 hrs, and live/dead cells measured by flow cytometry and trypan blue analysis. As expected, we observed 3 µM CDDP caused a 50% of mortality in A2870 cells non treated with sEVs (non-sEVs) (Fig. 6H and 6I). As previously, the sEVs fraction isolated in the presence of CDDP (chemo-sEVs) confers CDDP-cell death resistance to A2780 cells (Fig. 6H and 6I). Importantly, this cell death resistance phenotype was reverted when A2780 cells were tested in the presence of sEVs isolated in the presence of CDDP and rapamycin (Fig. 6H and 6I). Our findings indicate that the transfer of chemo-resistance to A2780 chemo-sensitive cells by chemo-sEVs secreted by A2780cis cells can be reduced by promoting lysosomal function through rapamycin mTORC1 inhibition.

## Discussion

Chemo-sEVs in OvCa are key players in the acquisition and transference of CDDP- chemoresistance (Safaei et al., 2005; Xavier et al., 2022; Yáñez-Mó et al., 2015). Therefore, the design of new strategies against OvCa and the development of chemoresistance to CDDP requires a deep understanding of the mechanisms related with the role of these sEVs in this type of cancer. In general, the type and quantity of EVs are controlled intracellularly through its biogenesis at the level of MVEs and ILVs and/or with the trafficking and fusion of MVEs with the PM (Steinbichler et al., 2019a; van Niel et al., 2018). In this work, we revealed that the A2780cis a cellular model of OvCa, that develops chemoresistance to CDDP, has an increased size of MVEs and increased number of ILVs. Consistent with this, A2780cis cells have high levels of ESCRTs and RAB27A proteins, which mediate ILVs biogenesis and secretion in A2780cis cells. Interestingly, there are transcriptional and post-translational pathways that could explain this phenotype. For example, NF-kB pathway increases the expression of RAB27A through RelA- p60, a mechanism that has been involved in CDDP chemoresistance in bladder cancer (Kan et al., 2020; J. Liu et al., 2017), however, it is unknown whether this mechanism could explain the RAB27A levels in A2780cis cells or in OvCa models. Additionally, the transcription factor EGR1, which controls RAB27A expression in other cellular models, is overexpressed in A2780cis cells (Ma et al., 2020; Rouillard et al., 2016). Moreover, high levels of RAB27A protein could also be the result of an increased protein stability related to its interaction with KIBRA, a scaffold protein involved in membrane trafficking such as the transport of early endosomes (Traer et al., 2007) in MVEs, which prevents RAB27A proteasomal degradation (Song et al., 2019). Although the link between CDDP and ESCRTs has been less investigated, it is known that CDDP promotes the activation of EGFR (Benhar et al., 2002), a growth factor known to induce the biogenesis of ILVs and the recruitment of ESCRTs to MVEs (Quinney et al., 2019). Altogether, these antecedents suggest that chemoresistance to CDPP in A2780cis cells might be the result of the activation in all these types of pathways, aspects that need further exploration.

Several RAB GTPases play a crucial role in the control of the trafficking of MVEs (Zerial & McBride, 2001). Among all RABs, we discovered A2780cis cells have high protein levels of RAB27A compared to A2780 cells (Fig. 1I, and 1J). In this regard, we propose that the acquisition of CDDP resistance is the result of adaptive tumoral cellular processes related to the biogenesis of ILVs and their secretion due to the overexpression of ESCRT machinery (Fig. 1G and 1H) and RAB27A (Fig. 1I and 1J). Previous studies suggested that the C13 CDDP-resistant model of OvCa cells secreted more EVs than its sensitive counterpart cellular model (Safaei et al., 2005). However, we did not observe a difference in the basal secretion of sEVs when we compared A2780 and A2780cis cell lines, observing only an increase when A2780cis cells were treated with CDDP, corresponding to chemo-sEVs (Fig. 2). Different post-translational modifications (PTMs) regulate the localization and function of different RABs, including phosphorylation, palmitoylation, ubiquitination, among others (Homma et al., 2021; Shinde & Maddika, 2018). Based on previous reports (Kielbik et al., 2018; Nguyen et al., 2017; Shi et al., 2021) CDDP could induce PTMs on RAB27A or in its effectors, causing the activation of RAB27A to promote chemo- sEVs secretion. In the tumor microenvironment, the chemoresistant cell population can spread its resistance mechanisms to remnant-sensitive cells (Madden et al., 2020; E. Yang et al., 2020). Interestingly, our results indicate that chemo-sEVs secreted in response to CDDP in A2780cis cells (Fig 2D), act as a mediator of cell-cell communication, transferring to A2780 cell the ability to evade CDDP-induced cell death (Fig. 2E and 2F). The cell-cell communication supported by sEVs is associated with the transference of messenger RNAs (mRNAs), miRNAs, DNAs, lipids, and proteins (Steinbichler et al., 2019b). Our proteomic analysis shows that CDDP changes the content of chemo-sEVs (Fig. 3A and 3C), whereas many of the differentially overrepresented proteins found could contribute to the transfer of chemoresistance to sensitive cells. For example, CHEK2 (Chk2) is a protein involved in DNA repair which is activated when DNA undergoes double strand break (DSB), promoting the arrest of the cell cycle (van Jaarsveld et al., 2020; J. Zhang et al., 2004). Similarly, we found XRCC6 (Ku70) and PRKDC, proteins involved in DSB repair by non-homologous end joining (Chen et al., 2021; Yu et al., 2018). Since CDDP induces DSB (Sears & Turchi, 2012; L. Wang et al., 2018), the transfer of these proteins by chemo-sEVs in response to CDDP could promote an activation of the DNA Damage response (DDR) in the recipient cells increasing its capacity to repair DNA, and therefore protecting sensitive cells against the DNA damage produced by CDDP. Supporting this idea, it has been reported that exosomes isolated from bone marrow mesenchymal stem cells (exoBM-MSCs) accelerated the DNA damage repair of recipient BM-MSCs treated with radiation, reducing the deleterious effect of this treatment (Zuo et al., 2020). Stabilization of p53 also plays an important role in DNA repair participating in the arrest of the cell cycle, thus giving time to the DNA repair machinery to facilitate genome stability (Williams & Schumacher, 2016). p53 is stabilized by the ubiquitination-deubiquitination cycle (Lee & Gu, 2009; M. Li et al., 2002). In this context, we observed that several proteins found enriched in the chemo-sEVs could be implicated in the balance between ubiquitination and deubiquitination of p53 (Fig. 3D). For example, the proteasomal deubiquitinase PSMD14, together with PSMD7 as a heterodimer, promotes p53 protein stability through a non-canonical pathway independent of the catalytic barrel of the 20S proteasome, function recently implicated in cancer progression (Bustamante et al., 2023). Interestingly, we found an enrichment of PSMD14 and PSMD7 in the chemo-sEVs isolated from A2780cis cells in response to CDDP. Therefore, it could be interesting to explore if this proteasome heterodimer could promote the stabilization of p53 in the recipient sensitive A2780 cells. Other possible proteins implicated could be related to the metabolism of the RNA such as ribosomal proteins (RPs), which have been suggested to play a role in drug resistance and cancer progression (Avitabile et al., 2022; Z. Huang et al., 2019; Kobayashi et al., 2006; C. Liu et al., 2019; Zhao et al., 2021). In fact, it has been reported in a model of CDDP-resistant gastric cancer that exosomes derived from these cells transfer chemoresistance to its sensitive counterpart via exosomal RPS3 (Sun et al., 2021). In this context, we found an enrichment of several RPs proteins, including RPS3, in the chemo-sEVs isolated from A2780cis cells in response to CDDP (Fig. 3D). Further investigation will help to define the more determinant players responsible for chemoresistance in sensitive cells. Proteomic analysis of chemo-sEVs is a nice strategy to get clues about these processes, however, the next challenge is to demonstrate which of these components contribute to cancer progression during chemotherapy.

In this study, we have underscored a fine regulatory mechanism that controls secretion of chemo-sEVs related with a step that defines the fusion of MVEs with the PM for secretion or alternatively with lysosomes for degradation, where the molecular details that exactly defines one or another direction are only partially understood (Huotari & Helenius, 2011; van Niel et al., 2018). Recently, it has been reported that the exchange between RAB7 and RAB27A on MVEs that controls exosome secretion is modulated by the ER membrane contact sites with endosomes (Verweij et al., 2022), opening a new hypothesis about how the secretion of chemo- sEVs is regulated. In this context, the levels of RAB27 and RAB7 in A2780cis cells seem to be mutually regulated, whereas upregulation of RAB27A correlates with a downregulation of RAB7, a phenotype that promotes the secretion of chemo-sEVs in response to CDDP (Fig 1I, J; 4M, N). Interestingly, hypoxia has been shown to trigger a similar phenotype regarding the balance of RAB27A/RAB7, a condition that also promotes an increased secretion of exosomes in chemoresistant ovarian cancer (Dorayappan et al., 2018). Regarding these findings, it is interesting the hypothesis that claims that the trafficking of MVEs is regulated by a fine balance between the status of lysosomal function and the cellular capacity for exosomal secretion (Eitan et al., 2016; Xu et al., 2022). Surprisingly, A2780cis showed a poor number of lysosomes, and the remnant lysosomes present had low acidity to have an appropriated degradative function (Fig. 4A-L). In agreement with previous findings (Kalayda et al., 2008; Safaei et al., 2005), we propose that the decay in lysosomal function together with an increase in key proteins needed for secretion, such as RAB27A, is mandatory to promote the biogenesis of MVEs and secretion of ILVs in response to CDDP in A2780cis cells (Fig. 2D), initial events in the development of chemoresistance. Additionally, the downregulation of RAB7 (Fig. 4M-N) might act as a key step to initiate the dysfunction of the lysosomal pathway in response to CDDP, an event that could decrease the delivery of hydrolases or proteins needed for pH maintenance (Saftig & Klumperman, 2009; Trivedi et al., 2020; C. Yang & Wang, 2021). In this context, it will be interesting to explore the impact of CDDP in the levels of those accessory proteins of AP-1 and GGAs complexes implicated in the delivery of hydrolases to lysosomes such as p56 (Mardones et al., 2007) and cyclin G-associated kinase (GAK) (Kametaka et al., 2007), among several others.

On the other hand, lysosomes have a drug-sequestering function that diminish the availability of the drugs within cells including CDDP (Galluzzi et al., 2014; Geisslinger et al., 2020; Zhai & el Hiani, 2020; Zhitomirsky & Assaraf, 2016). Thus, the process of chemotherapy resistance may be driven by the sequestration of CDDP within lysosomes, leading to lysosomal dysfunction in A2780cis cells during the early stages of resistance acquisition (Fig. 4). This sequestration can cause the secretion of sEVs that can become more pronounced over time due to high levels of RAB27A, resulting in the orchestration of a cell survival program and the development of chemotherapy resistance. Thus, we propose RAB27A-dependent sEVs secretion is a crucial pathway activated under lysosomal dysfunction and a critical feature for CDDP chemoresistance in OvCa.

In addition to the direct role of the CDDP in lysosomes it is possible that CDDP could activate a signaling pathway with negative impact on lysosomes. In this regard, it has been demonstrated that CDDP activates mTORC1 causing the inactivation of the transcription factor TFEB (O’Dea et al., 1988; Palmieri et al., 2011). Consistently, CDDP in A2780cis cells activates mTORC1 (Fig. 6A and 6B), a phenotype that could be responsible for lysosomal dysfunction (Fang et al., 2022). Additionally, A2780cis cells show a reduction of RAB7 protein levels, suggesting the involvement of this GTPase in the decline of the lysosomal function during the acquisition of chemoresistance.

Importantly, the silencing of RAB27A in A2780cis cells, a strategy that inhibits the secretion of exosomes in a variety of other cellular models (Blanc & Vidal, 2018; Bobrie et al., 2012; H. Huang et al., 2021; Ostrowski et al., 2010; Salimu et al., 2017), promotes an increase in the protein levels of RAB7, confirming a closed regulation between these two GTPases. Moreover, silencing of RAB27A also causes a recovery in the number and function of lysosomes, together with a decrease in CD63 strongly indicates that silencing of RAB27A enhances the delivery of content into the endo-lysosomal degradative pathway. Further studies should be addressed if overexpression of RAB7 could be sufficient to recapitulate the effect observed with the silencing of RAB27. Moreover, we found that inhibition of mTORC1 with rapamycin, a well-known strategy to promote lysosomal biogenesis, affects the secretion of chemo-sEVs in A2780cis cells in response to CDDP. Moreover, the chemo-sEVs secreted have a reduced capacity to transfer chemoresistance in sensitive cells. Altogether, these findings confirm a close interplay between the lysosomal function and the capacity of cells to secrete chemo-sEVs, in which the interruption of one pathway impacts positively the other.

These findings offer possible targets of intervention for OvCa patients that develop chemoresistance to CDDP including drugs with a potent inhibitory effect on sEVs secretion such as specific inhibitors of RAB27A activity (H. Zhang et al., 2020). Another strategy could be related with an induction of the lysosomal function through the biogenesis of these organelles with the use of rapamycin, a compound that enhances the anti-tumoral effects of derivatives of CDDP (oxaliplatin) in A2780cis (J. Liu et al., 2015). Moreover, high levels of RAB7 in C13 CDDP-resistant model of OvCa cells has been demonstrated to sensitize cells to CDDP (Guerra et al., 2019), that based on our results could be due to an increase in the degradative endo- lysosomal pathway and lysosome biogenesis.

The results of this study provide insight into the development of CDDP chemoresistance in OvCa. By uncovering the interplay between lysosomal function and the secretion of chemo-sEVs, two new potential strategies against CDDP chemoresistance were identified: silencing of RAB27A and inhibition of mTORC1 with rapamycin. These interventions aim to revert the lysosomal dysfunction found in CDDP-resistant cancer cells and may improve the efficiency of CDDP-based cancer treatments. The findings of this study have the potential to inform the design of complementary strategies that incorporate lysosomal-targeting agents to avoid CDDP chemoresistance.

## Legends Graphical Abstract

A2780 CDDP-sensitive cells are derived from an epithelial ovarian tumor. A2780cis CDDP-resistant cells were generated by the exposure of A2780 cells at high concentrations of CDDP for 6 months (*1*). A2780cis CDDP-resistant cells have more machinery involved in ILVs/sEVs biogenesis (*2*) and capacity to secrete sEVs (*3*) than A2780 CDDP-sensitive cells. The secretion of exosomes by A2780cis cells only increases in response to CDDP, and these have resistant transference capacity (*4*). This phenotype of A2780cis cells is associated with a lysosomal alteration (reduced number of lysosomes, with less acidity and degradative activity) and reduced endolysosomal pathway (*5*). The absence of RAB27A in A2780cis CDDP-resistant cells reverts the lysosome phenotype and endolysosomal pathway, such as A2780 CDDP-sensitive cells. Similarly, increasing lysosomal biogenesis in A2780cis CDDP-resistant cells by rapamycin treatment prevents the effect of exosome secretion and their resistance transference capacity induced by CDDP (*6*).

---

Figure S1. RAB27A KD recovers the lysosomal acidity and degradative function in A2780cis CDDP-resistant OvCa cells. **(A)** Confocal live cell images of A2780 shLuc, A2780cis shLuc, and A2780cis shRAB27A cells were incubated with LysoTracker Red and HOECHST. Scale Bar 10 μm. **(B-C)** Analysis of number (B) and average intensity (C) of LysoTracker Red structures from images as those shown in (A). Bars indicated the mean with SEM; ***P<0.001; one-tiled, unpaired, parametric t-Test (n>50 cells). **(D)** Confocal live cell images of A2780 shLuc, A2780cis shLuc, and A2780cis shRAB27A cells were incubated with Magic Red and HOECHST33342. Scale Bar 10 μm. **(E)** Analysis of average intensity of Magic Red structures from images as those shown in (E). Bars indicated the mean with SEM; ***P<0.001; one- tiled, unpaired, parametric, t-Test (n>50 cells).

