## Supplementary figures and images for "Chemo-sEVs release in cisplatin-resistance ovarian cancer cells are regulated by the lysosomal function"

### Supplemental Figure 1

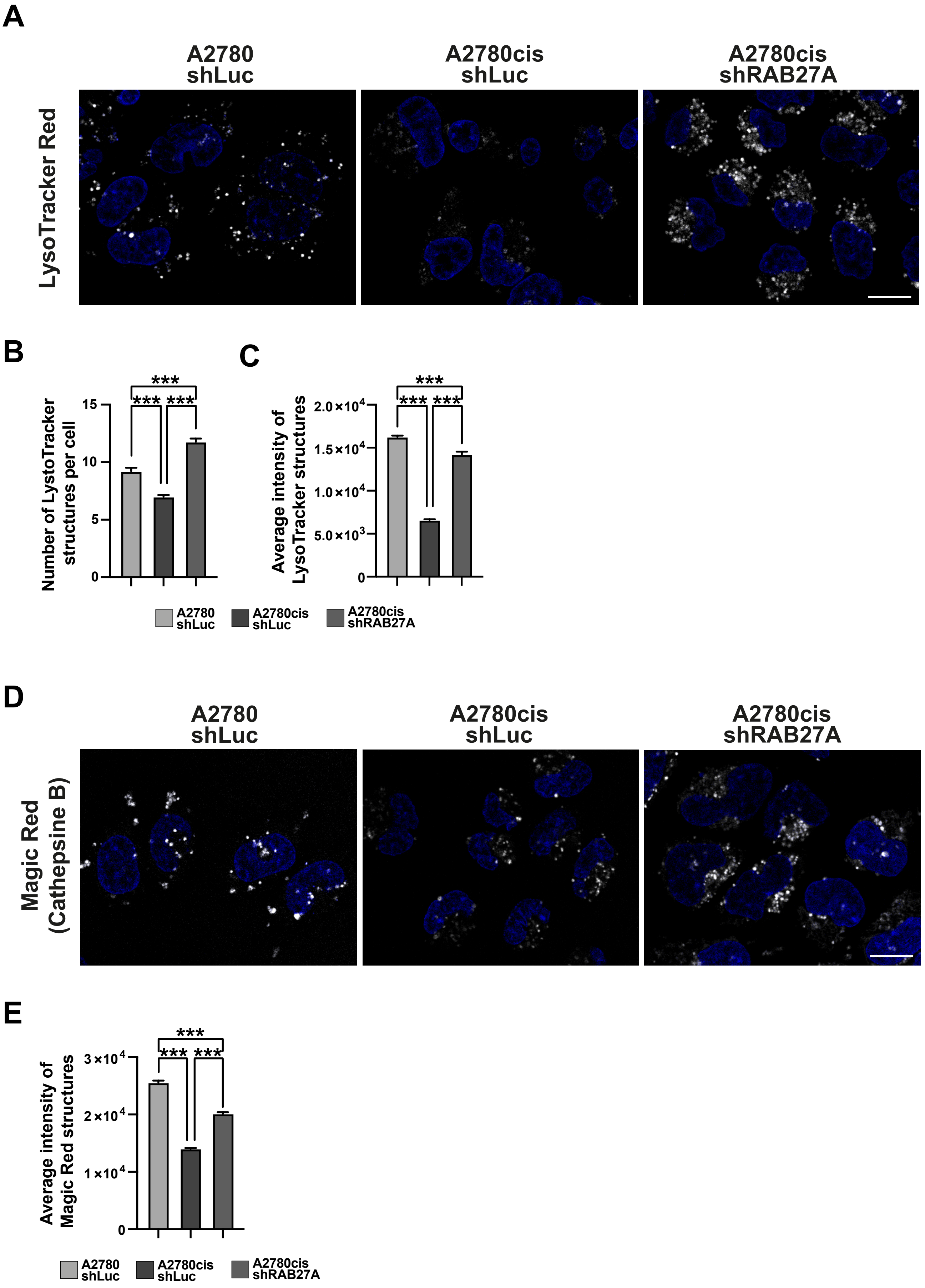


*Fig S1. Cerda-Troncoso et al.*
